# Circulating miRNA diversity, origin and response to changing metabolic and reproductive states, new insights from the rainbow trout

**DOI:** 10.1101/2021.04.27.440972

**Authors:** E Emilie Cardona, C Cervin Guyomar, Thomas Desvignes, J Jérôme Montfort, Samia Guendouz, John H. Postlethwait, S Sandrine Skiba-Cassy, Julien Bobe

**Author notes:** equal contribution (E Emilie Cardona, C Cervin Guyomar).

## Abstract

Circulating miRNAs (c-miRNAs) are found in most, if not all, biological fluids and are becoming well established biomarkers of many human pathologies. The aim of the present study was to investigate the potential of c-miRNAs as biomarkers of reproductive and metabolic states in fish, a question that has received little attention. Plasma was collected throughout the reproductive cycle from rainbow trout females subjected to two different feeding levels to trigger contrasting metabolic states; ovarian fluid was sample at ovulation. Fluid samples were subjected to small RNA-seq analysis followed by quantitative PCR validation for a subset of promising c-miRNA biomarkers. A comprehensive miRNA repertoire, which was lacking in trout, was first established to allow subsequent analysis. We first showed that biological fluids miRNAomes are complex and encompass a high proportion of the overall species miRNAome. While sharing a high proportion of common miRNAs, plasma and ovarian fluid miRNAomes exhibited strong fluid-specific signatures. We further showed that the plasma miRNAome exhibited major significant changes depending on metabolic and reproductive state. We subsequently identified three (miR-1-1/2-3p, miR-133-a-1/2-3p and miR-206-3p) evolutionarily conserved muscle-specific miRNA that accumulate in the plasma in response to high feeding rates, making these myomiRs strong candidate biomarkers of active myogenesis. We also identified miR-202-5p as a candidate biomarker for reproductive success that could be used to predict ovulation and/or egg quality. These highly promising results reveal the high potential of c-miRNAs as physiologically relevant biomarkers and pave the way for the use of c-miRNAs for non-invasive phenotyping in various fish species.

## 1 Introduction

MicroRNAs (miRNAs) are small non-coding RNAs (about 22nt in length) that act as post-transcriptional gene regulators by inducing mRNA decay or translational repression in animals and plants [1,2]. In vertebrates, miRNAs are highly conserved and associated with numerous physiological and pathological processes [3]. miRNAs can be secreted from cells into body fluids, such as serum, plasma, saliva, colostrum, milk, urine, semen, amniotic fluid, cerebrospinal fluid, peritoneal fluid, and pleural fluid [4–7], where they remain particularly stable [8]. In the past decade, microRNAs have emerged as highly promising biomarker molecules, due to their presence and stability in most biological fluids, including blood plasma. Circulating miRNAs (c-miRNAs) have been documented in many biomedical contexts and have revealed great potential as diagnostic and prognostic biomarkers in human medicine for a variety of pathologies including cancer [9]. In contrast, c-miRNAs in non-pathological contexts have received little attention and data in non-human species remain scarce and still for the most part related to pathological issues [10–13]. In a few cases, c-miRNAs have, however, been studied in non-pathological contexts, such as puberty [14], oestrus cycle and pregnancy [15,16], and embryonic development [17] but also in relation to animal nutrition [18–20], or in response to environmental changes [21–24].

To date, very little data exist on c-miRNAs in aquatic species, including in fish, even though the presence of miRNAs was recently reported in the plasma of Senegalese sole and rainbow trout [20,25], in the seminal fluid of Atlantic salmon [26] and in the mucus of rainbow trout [25]. In order to monitor the growth, reproduction, and health of stocks in aquaculture or of a wild fish population, current methods necessitate important, repeated, and often stressful manipulations of the fish to collect live measurements. Furthermore, numerous representative specimens often need to be euthanized to assess the reproductive state for example. In contrast, non-invasive sampling of body fluids such as plasma could potentially inform not only of the fish present condition but also its past and present overall growth, reproductive advancement, and health status. Therefore, developing c-miRNAs that could serve as biomarkers of physiological, metabolic, and health status would be of major interest for aquaculture but also for wild population management, including endangered species, for which each wild and captive specimen is critical.

The aim of the present study was thus to investigate the potential of c-miRNAs as biomarkers of reproductive and metabolic states. We thus aimed at characterizing the rainbow trout c-miRNAome in two biological fluids: plasma and ovarian fluid (i.e., the fluid in which the eggs are bathed after ovulation), which can be easily and non-invasively collected. We investigated two different feeding regimes that trigger contrasting metabolic states [27]. Plasma was analyzed at different stages of the reproductive cycle and ovarian fluid was studied at ovulation. Rainbow trout was chosen due to its relatively large size, which simplified fluid collection, as well as its sequenced genome and available small RNA-seq data in 38 different tissues, cell types, organs, and embryonic stages [28]. In fish, a comprehensive view of the overall miRNA repertoire was only recently characterized [29–31] but was still lacking in rainbow trout. We thus established a comprehensive annotation of expressed miRNAs in rainbow trout using existing and newly generated sRNA-seq data and using existing miRNAome annotations in teleost fish [29–33]. To gain insight into the organ of origin, diversity and specificity of c-miRNAomes, plasma and ovarian fluid miRNA expression data were analyzed along with data from a panel of 21 tissues and organs. We then evaluated the potential of c-miRNAs to serve as biomarkers of the reproductive stage of female rainbow trout throughout their reproductive cycle. Finally, we evaluated the potential of selected c-miRNAs to serve as biomarkers of the metabolic state through the use of two feeding levels: *ad libitum* feeding or a physiologically relevant moderate feeding restriction. We used a stepwise strategy, relying first on the small RNA sequencing of a limited number of pools of biological samples to identify major c-miRNAs and then further validating their potential using quantitative PCR on a large number of biological replicates and additional reproductive stages. Here we show that plasma and ovarian fluid miRNAomes are complex and share a majority of common miRNAs. We report, however, strong fluid-specific signatures both in terms of expression levels and specific miRNAs and identify candidate biomarkers of sexual maturation and active myogenesis. Together, these results highlight the relevance of c-miRNAs for non-invasive phenotyping in aquatic species.

## 2 Results and discussion

### 2.1 The rainbow trout reference miRNAome

In order to evaluate the complexity of the rainbow trout circulating miRNAome, we first had to establish a comprehensive rainbow trout miRNAome that could be used as a reference. Using a total of 52 sequencing libraries composed of 14 new libraries (plasma and ovarian fluid) and 38 libraries that we had previously generated in a wide variety of tissues, cellular populations or organs, including whole embryo [28], we were able to identify and annotate 354 mature rainbow trout miRNAs corresponding to at least 280 miRNA genes adapting a strategy previously used in different fish species [29,31]. This represents, to our knowledge, the first reference rainbow trout miRNA repertoire annotation. While slightly lower, the number of mature rainbow trout miRNAs reported here is consistent with previous reports in other ray-finned fish species using the same strategy that led to the annotation of 362, 495, 396, and 408 individual mature miRNAs in gar, zebrafish, stickleback, and blackfin icefish, respectively [29–31]. Nevertheless, the present report provides a comprehensive and evolutionarily-supported rainbow trout miRNA repertoire annotation, that was previously missing in this species.

### 2.2 The high complexity of circulating miRNAome

Among the 354 annotated rainbow trout miRNAs, 331 were detected above a threshold of 10 reads per million reads, on average, either in one of two biological fluids studied or in one of the 21 other types of analyzed samples (brain, eggs, gills, gonad, head-kidney, heart, intestine, leucocytes, liver, muscle, myoblasts, myotubes, ovary, pituitary, skin, spermatogonia, spleen, stomach, testis, trunk-kidney, whole embryos). In biological fluids, 211 miRNAs could be identified that corresponded to 64% of the overall expressed miRNAome diversity (Figure 1A). Among these 211 miRNAs, 172 (82%) were detected in both plasma and ovarian fluid, while 24 (11%) were detected only in plasma and 15 (7%) appeared only in ovarian fluid (Figure 1A). Notably, two miRNAs (miR-365-2-5p, miR-23b-2-5p) were detected above the 10 RPM threshold only in plasma but not in any other studied sample, while a single miRNA (miR-726-5p) was only found in ovarian fluid (Figure 1A). A comprehensive analysis of miRNA expression levels revealed that the overall distribution patterns of miRNA read counts were similar in biological fluids and in other samples (Figure 1B). In each analyzed library, including ovarian fluid and plasma libraries, a few miRNAs accounted for most of the reads per million (RPM). Together these data illustrate the relatively large complexity of the c-miRNA repertoire present in the two biological fluids studied here. Our results are consistent with existing data in humans, chicken and cow plasma in which 349, 649, and 468 miRNAs were reported, respectively [4,21,24]. This trout result, however, is to our knowledge the first comprehensive characterization of plasma and ovarian fluid miRNAomes in a fish species.

**Figure 1:**
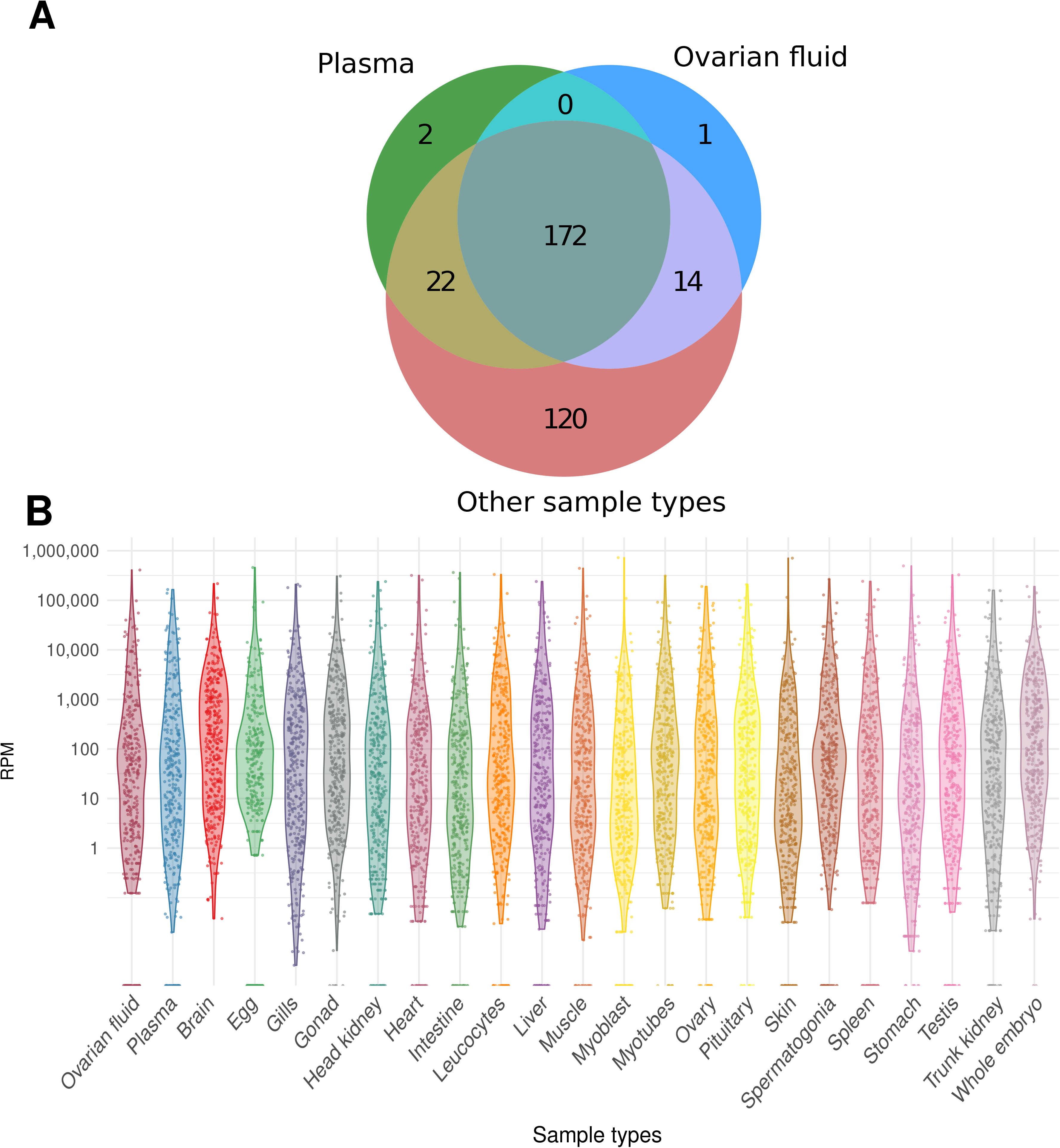
Circulating miRNA repertoires. A: Venn diagram of miRNAs detected in plasma, ovarian fluid and in 21 other types of samples (brain, eggs, gills, gonad, head-kidney, heart, intestine, leucocytes, liver, muscle, myoblasts, myotubes, ovary, pituitary, skin, spermatogonia, spleen, stomach, testis, trunk-kidney, whole embryos). A miRNA was considered expressed in a type of samples when its normalized abundance exceeded 10 RPM (reads per million reads), on average. B: Distribution of miRNA normalized counts across types of samples listed above in RPM.

### 2.3 Origin and specificity of c-miRNA in plasma and ovarian fluid

To investigate the possible organ of origin of miRNAs present in plasma and ovarian fluid, we categorized c-miRNAs based on the organ in which they exhibited the highest expression, under the hypothesis that an organ strongly expressing a miRNA is likely an organ secreting the miRNA into the fluid, or at least one of the major contributors. This analysis investigated a limited subset of 13 different organs (brain, gills, heart, head kidney, intestine, liver, muscle, ovary, pituitary, skin, spleen, stomach, trunk kidney) in order to avoid male tissues and organs, complex libraries (e.g., whole embryos), individual cell types (e.g., myoblasts). We observed that miRNAs present in both ovarian fluid and plasma had maximum expression (i.e., were detected at the highest level) in a wide diversity of organs (Figure 2A). For both analyzed fluids, brain, gills, pituitary, ovary, and liver were the organs in which miRNAs had a maximum expression and no major difference in potential organs of origin could be identified between these two fluids (Figure 2A). These data suggest that many organs might contribute to the complexity of c-miRNAome in both plasma and ovarian fluid.

**Figure 2:**
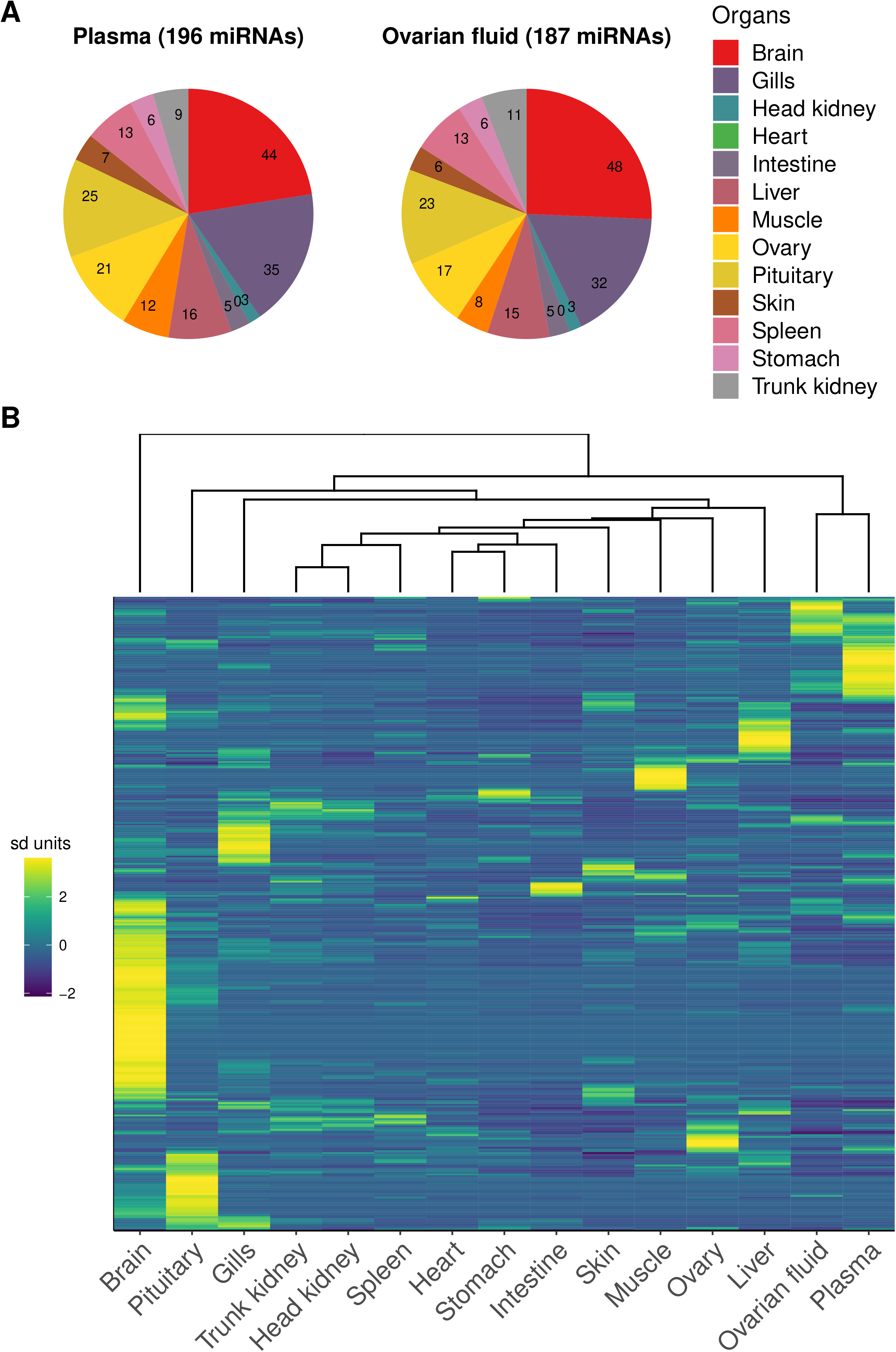
Origin and expression patterns of c-miRNAs. A: For each fluid c-miRNA, the organ in which its expression was the highest was considered as a possible organ of origin. The number of c-miRNAs exhibiting highest expression in a specific organ are shown on the chart. Heart is displayed even though no c-miRNA had a maximum expression in this organ. B: Heatmap of miRNA expression across different organs, plasma and ovarian fluid. The heatmap was built using a row-scaled matrix of the normalized read counts (RPM) supplied by *Prost!*. Normalized RPM counts were averaged for all samples of a given sample type.

When miRNA expression in fluids was analyzed together with expression data in a panel of 13 organs from which they could originate, we observed that both plasma and ovarian fluid exhibited specific expression patterns as distinct as in the other analyzed organs, if not more. The heatmap presented in Figure 2B clearly shows that, while sharing common strongly expressed miRNAs, each fluid is nonetheless characterized by the overabundance of several miRNAs (the yellow ones towards the top right of the panel) that are not overabundant in any other part (organ or tissue) of the body analyzed in the present study (Additional file 1). The PCA analysis carried out using all available samples clearly illustrated that biological fluid miRNAomes, while distinct, are also clearly different from all other “solid” tissue/organ miRNAomes (Figure 3A). When analyzing the presence of miRNAs in the different libraries, we observed that most miRNAs present in plasma and ovarian fluid were also detected in most organs (Figure 3B).

**Figure 3:**
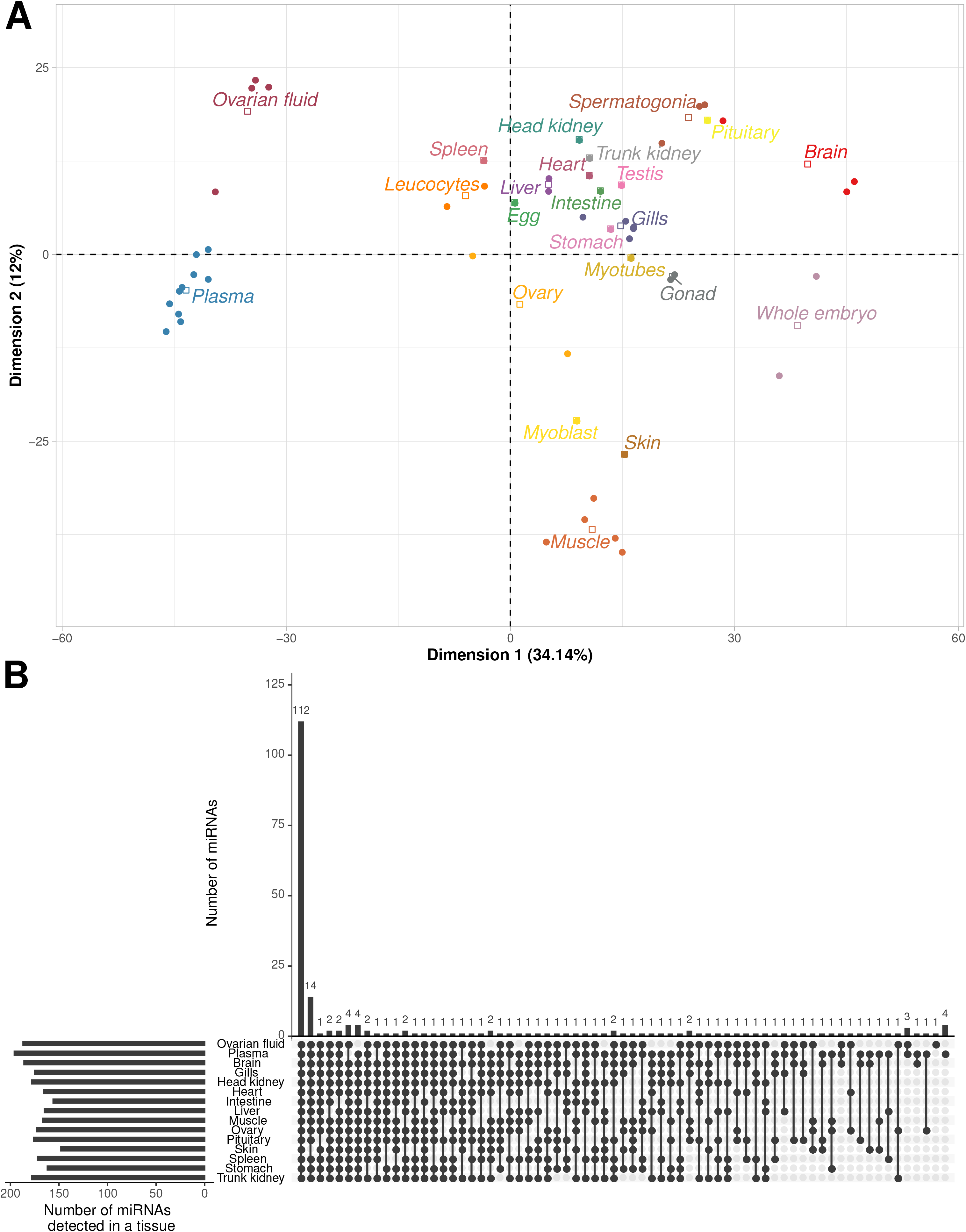
Detection of plasma and ovarian fluid miRNAs in various organs. A: PCA analysis of normalized RPM supplied by Prost! using all samples (N=52). PCA was centered but not scaled. B: The number of circulating miRNAs detected in all possible combinations of analyzed libraries is displayed. Libraries in which miRNAs were detected are indicated by a black dot. miRNAs were considered present in an organ or a fluid if their normalized abundance exceeded 10 RPM, on average, for each type of sample. The analysis was limited to the two biological fluids (plasma and ovarian fluid) and a subset of 13 different organs displayed in figure 2.

When analyzing the expression of the miRNAs exhibiting strong expression in ovarian fluid and plasma, we observed that among the 10 most abundant miRNAs in ovarian fluid and plasma, seven (miR-451-5p, let-7a-5p, miR-21-5p, miR-16b-5p, miR-26a-5p, let-7e-5p, miR-30d-5p) were common to both fluids. In plasma miR-92a-3p, miR-150-5p, and miR-128-3p were the three other most abundant miRNAs. In ovarian fluid, miR-202-5p, miR-22a-1-3p, and miR-146a-5p were the three other most abundant miRNAs. For miR-451-5p, miR-16b-5p, miR-26a-5p, and miR-92a-3p, a clear over abundance was observed in both fluids in comparison to body organs (Figure 4). In contrast, the other most abundant miRNAs in fluids were also highly abundant in at least one other organs (Figure 4).

**Figure 4:**
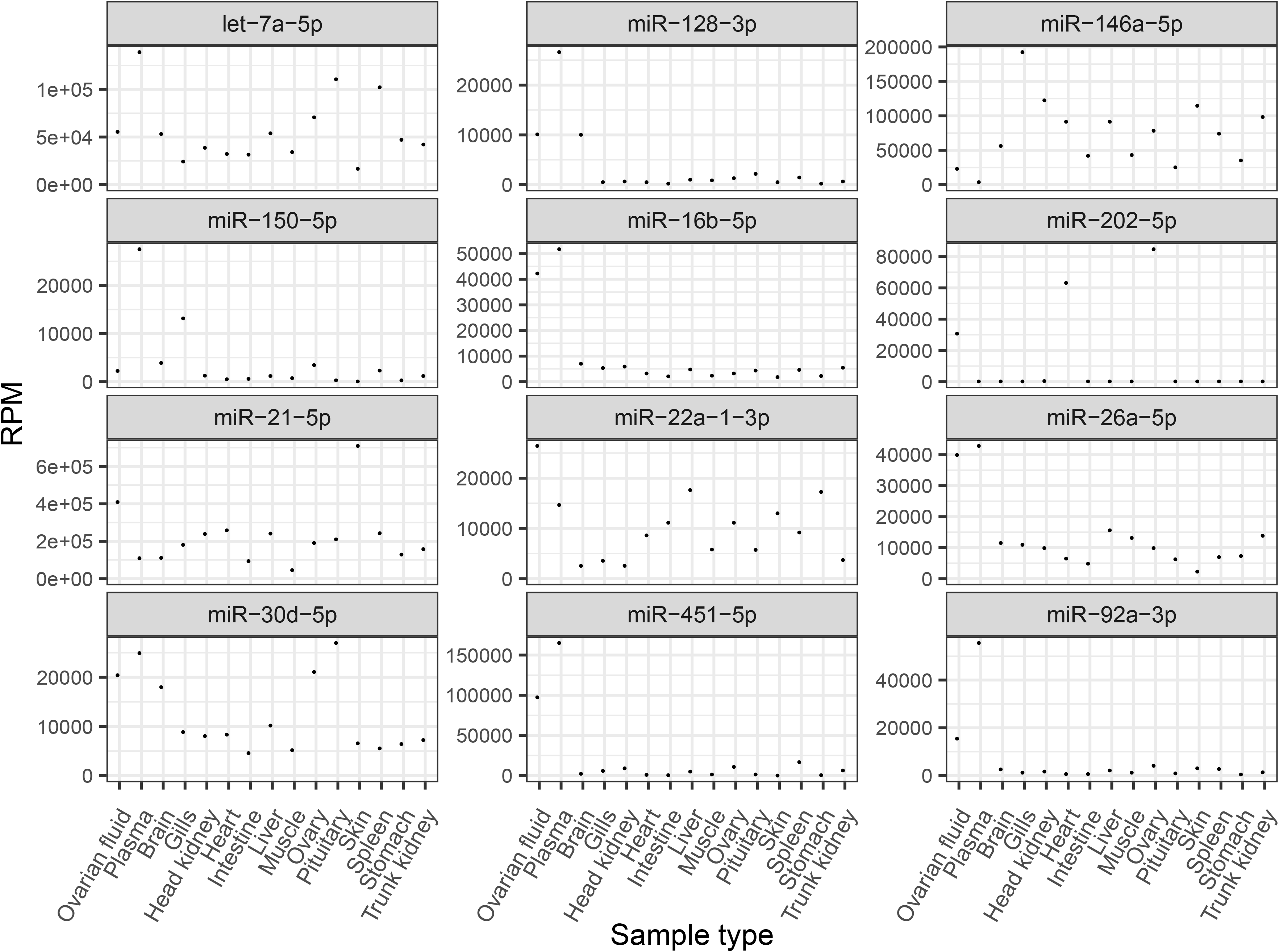
Normalized expression values of the most abundant c-miRNAs across organs and biological fluids. The analysis was performed using the two biological fluids (plasma and ovarian fluid) and a subset of 13 different organs displayed in figure 2. Normalized RPM counts were averaged for all samples of a given sample type. miR-451-5p, let-7a-5p, miR-21-5p, miR-16b-5p, miR-30d-5p, and miR-26a-5p were among the 10 most abundant miRNAs for both plasma and ovarian fluid. miR-92a-3p, miR-150-5p and miR-128-3p were among the most abundant miRNAs in plasma. miR-202-5p, miR-22a-1-3p and miR-146a-5p were among the most abundant miRNAs in ovarian fluid.

Together, these data indicate that c-miRNA repertoires in rainbow trout are complex. Similar to what is observed for miRNAs in different organs, c-miRNAomes in plasma and ovarian fluid each exhibit specific expression profiles, with fluid-specific combinations of highly expressed miRNAs, major differences in miRNA abundances and fluid-type specific miRNAs in comparison to the other fluid and organs analyzed. These data suggest that the complexity of c-miRNAomes in plasma and ovarian fluid results, at least in part, from the accumulation of miRNAs originating from a wide diversity of organs.

The presence in body fluids of miRNAs that cannot be detected in other organs, or detected at much lower levels, suggests that these miRNAs originate from other sources that were not investigated here. These specific miRNAs could also originate from miRNA-expressing cells present in these biological fluids or in their vicinity. For example, miR451-5p could originate from erythrocytes that greatly express this miRNA during the late stage of red-blood cell maturation [34–37] or miR21-5p could be expressed by the endothelial cells forming the vasculature [38]. Finally, it is also possible that the level of these miRNAs results from their progressive accumulation in these fluids over time permitted by their high stability in nuclease-rich fluids [39].

### 2.4 Differences and similarities of ovarian fluid and plasma miRNA repertoires

As indicated above, plasma and ovarian fluid have many miRNAs in common. Marked differences between plasma and ovarian fluid miRNAomes, however, exist in terms of both miRNA profiles and expression of fluid-specific miRNAs. The PCA analysis of fluid samples only (Figure 5A) clearly shows that the overall c-miRNA profiles differ between plasma and ovarian fluid samples. Accordingly, 138 miRNAs were significantly differentially abundant between ovarian fluid and plasma (additional file 2). Among these 138 miRNAs, 67 were over-abundant in ovarian fluid and 71 in plasma.

**Figure 5:**
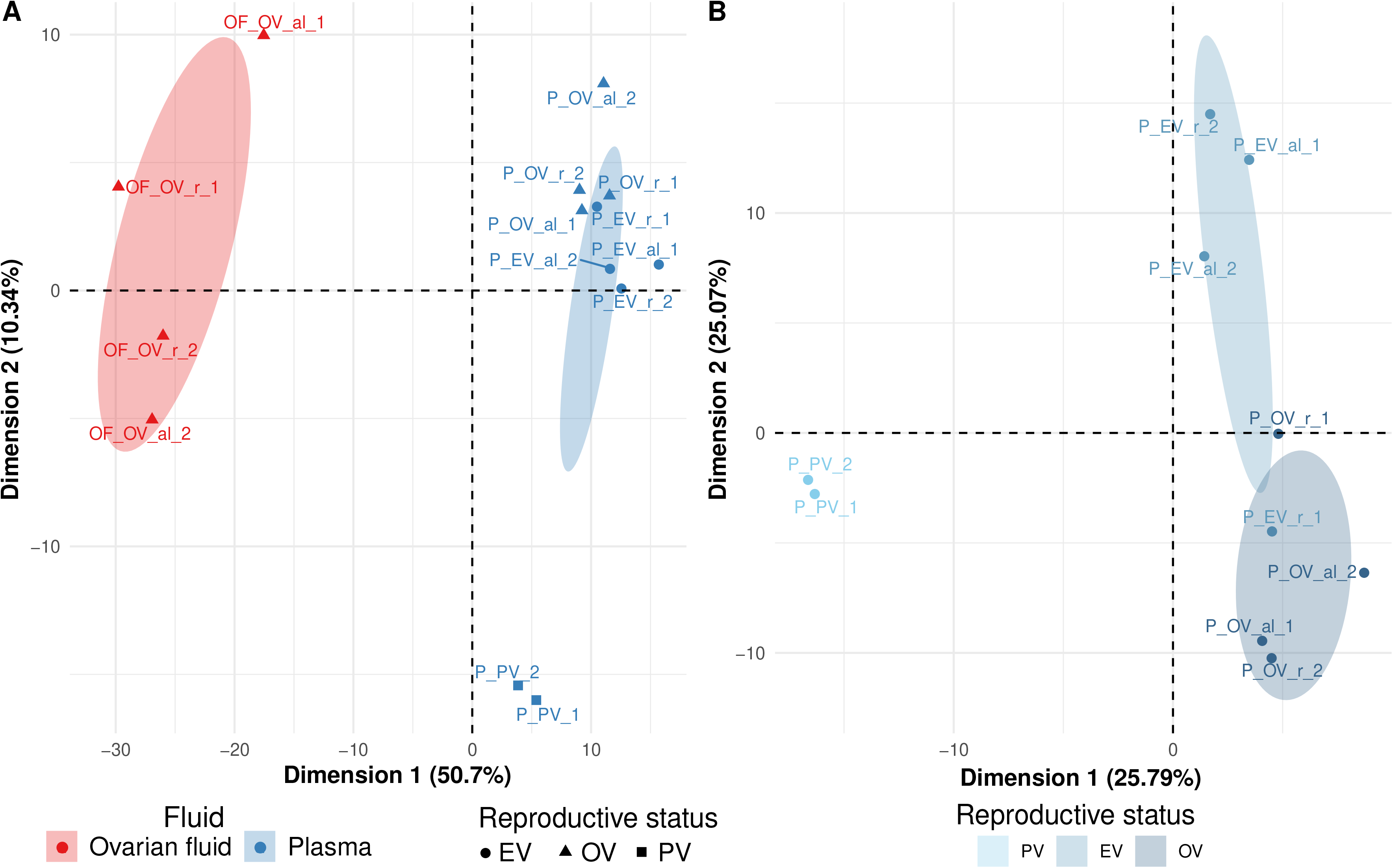
Principal Component Analyses of biological fluid c-miRNAs. PCAs were centered but not scaled and computed from normalized miRNAs counts (RPM) supplied by Prost!. Ellipses represent 95% confidence intervals, and were drawn for conditions represented by at least three individuals. Points are samples named under the following pattern: “Fluid_Reproductive stage_feeding_replicate”. The value for “Fluid” is either plasma (P) or ovarian fluid (OF). The following time points were analyzed: pre-vitellogenesis (PV), early vitellogenesis (EV), and ovulation (OV). The feeding level was either *ad libitum* (al) or restricted (r). A: PCA of all 10 plasma and 4 ovarian fluid samples. B: PCA using only the plasma samples (N=10) collected at the three different reproductive stages.

Our results are consistent with previous studies in humans showing distinct miRNA compositions in different fluid types [4]. While the authors [4] suggested a common origin for miRNAs present in the different body fluids, they also reported fluid-specific enrichment of several miRNAs including in plasma. In the present study, two miRNAs (miR-202-5p and miR-194b-5p) exhibited a dramatic, over 100-fold, enrichment in ovarian fluid in comparison to plasma (Additional file 2). In contrast, two miRNAs (miR-460-5p and miR-365-2-5p) exhibited the opposite pattern. While the function of extracellular miRNAs remains unclear [39] it has been hypothesized that fluid-specific miRNAs could have regulatory functions in surrounding tissues [4]. In rainbow trout, as in many vertebrates, miR-202-5p is predominantly expressed in gonads [28,40–44]. The strong abundance of miR-202-5p in ovarian fluid therefore agrees with its strong ovarian expression because the ovarian fluid, in which ovulated oocytes (i.e. unfertilized eggs) are held in the body cavity until spawning, is, at least in part, from ovarian origin [45]. Conversely, the plasma perfuses all organs and transports molecules throughout the body, including in the ovaries [4] and many ovarian fluid components such as proteins are also known to be brought in from the plasma [46,47]. Ovarian fluid c-miRNAs may thus also, in part, be brought in the ovarian fluid via the blood, which would be consistent with the presence of a high proportion of common miRNAs in both fluids. In medaka, miR-202-5p plays a major role in female reproduction, specifically in the control of egg production and egg ability to be fertilized [40]. The overabundance of miR-202-5p in ovarian fluid, compared to plasma, would be consistent with a physiological role of miR-202-5p in the ovarian fluid before, at, or after ovulation, a period associated with major events, including the final maturation of oocytes and the onset of the next reproductive cycle.

### 2.5 Varying c-miRNA abundance in response to reproductive and metabolic states

In the present study, we also aimed at identifying plasma c-miRNAs that change in expression level during the reproductive cycle or in response to different metabolic state resulting from different feeding levels. The PCA analysis (Figure 5B) revealed clear differences in plasma global c-miRNAomes during the reproductive cycle. Differences are especially noticeable between samples taken at the beginning of the reproductive cycle (i.e., at previtellogenic stage) vs. fish sampled later during the reproductive cycle. The statistical analysis led to the identification of 107 differentially abundant miRNAs during the reproductive cycle (Additional file 3). The heatmap of these miRNAs presented in Figure 6A revealed four different clusters of miRNA expression profiles during the reproductive cycle. Most changes in expression occurred between previtellogenesis (PV) and early-vitellogenesis (EV). The differential expression analysis resulted in the identification of 48 downregulated (cluster 1) and 50 upregulated (cluster 2) miRNAs in PV compared to EV (Figure 6B). In addition, we identified five miRNAs upregulated at ovulation (OV, cluster 3) and four miRNAs exhibiting the opposite pattern (cluster 4). Together, these results indicate that major changes occurred in the plasma c-miRNAome during the reproductive cycle and that a significant portion of the plasma c-miRNAome (107 c-miRNAs, 55% of the overall plasma c-miRNAome) exhibits a differential abundance between at least two stages of the reproductive cycle. These results show that the overall plasma miRNAome exhibits marked stage-specific signatures during the reproductive cycle.

**Figure 6:**
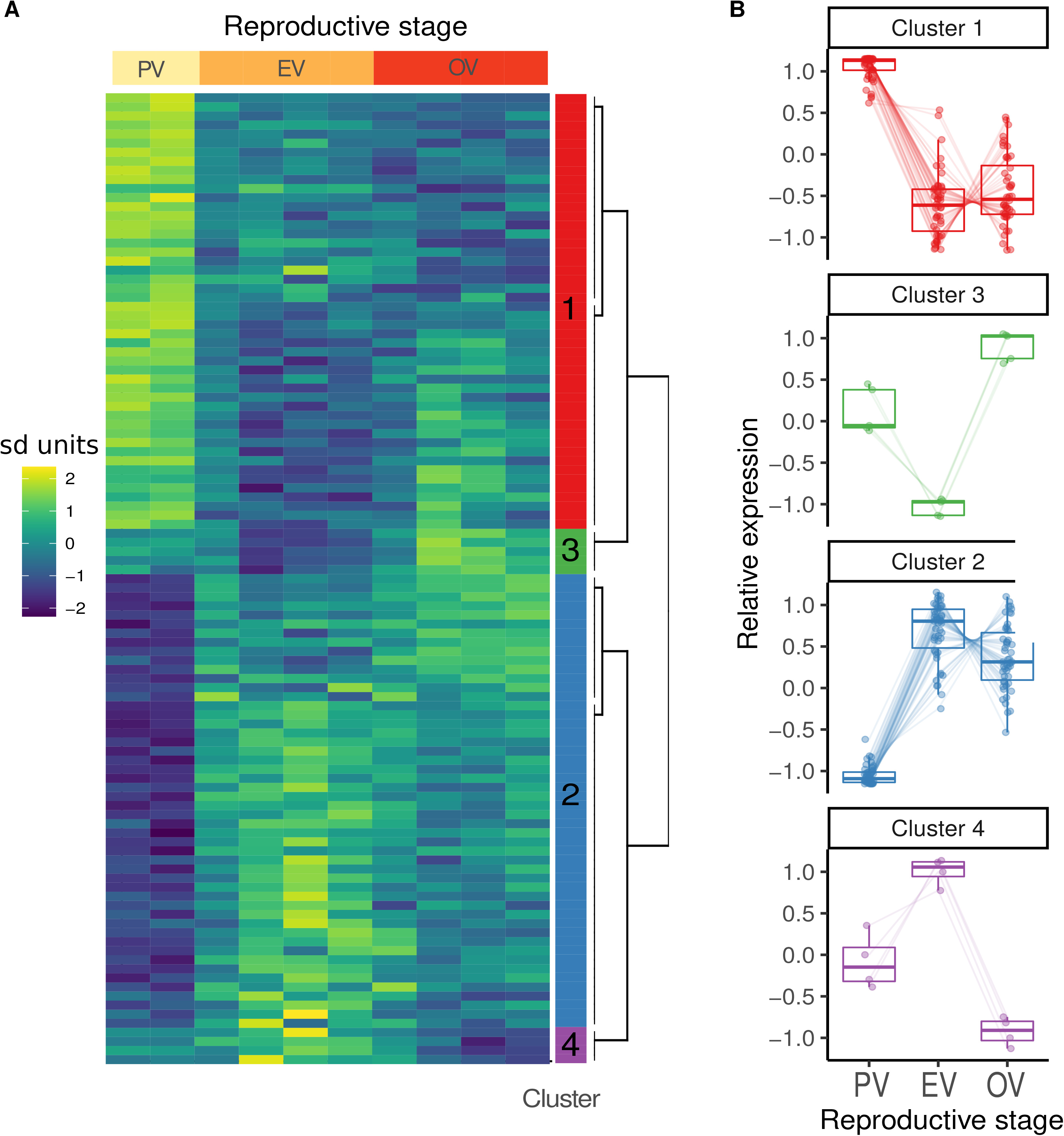
Expression profiling of c-miRNAs differentially expressed during the reproductive cycle. A: Expression heatmap for c-miRNAs differentially expressed across time. Expression values were log transformed read counts of the 107 c-miRNAs differentially expressed during the reproductive cycle. The heatmap was scaled by row. Expression dynamic clusters are indicated by colors in the rightmost column. B: Expression dynamics by cluster of the 107 differentially expressed c-miRNAs. All differentially expressed miRNAs were assigned to one of four clusters based on their expression dynamic along time.

In contrast, we were not able to detect any significant differences in c-miRNA abundances in the plasma in response to metabolic levels (i.e., feeding level) (“al” for *ad libitum* and “r” for restricted diet in Figure 6). This result could, however, result from the low number of replicates (two per diet) that composed our RNA-seq dataset.

### 2.6 Circulating plasma miRNAs as non-invasive biomarkers of metabolic and reproductive states

To evaluate the potential of plasma c-miRNAs to respond to differences in metabolic levels we selected the most promising miRNAs (i.e., exhibiting the highest fold-changes in our small RNA-seq data) and conducted an extended analysis of their expression by quantitative PCR (QPCR) using five replicates and an additional time point during the reproductive cycle (Late vitellogenesis, LV) (Figure 7). The potential origin of these candidate c-miRNAs was also analyzed by QPCR in a panel of organs to shed light on their possible origin of expression (Figure 8). Candidate c-miRNAs were selected based on their expression fold-change between different metabolic levels or reproductive stages to maximize their relevance as potential biomarkers. Quantitative PCR demonstrated that selected biomarker c-miRNAs exhibited highly significant changes in their plasma abundance throughout the reproductive cycle. In most cases, these changes occurred in a feeding level-dependent manner, indicating that circulating miRNA levels in plasma can be deeply influenced by metabolism. Among analyzed candidate biomarkers, miR-1-1/2-3p, miR-133-a-1/2-3p and miR-206-3p exhibited a similar pattern throughout the reproductive cycle with a dramatic increase in plasma abundance during vitellogenesis (i.e. the reproductive phase characterized by major yolk protein uptake from the blood stream by the oocyte) when fish were fed *ad libitum* but not when the food was restricted (Figure 7A-C). When investigating the organs expressing these three miRNAs, we observed a predominant expression in skeletal muscle (Figure 8). These miRNAs are known to be “muscle-specific” and often referred to as “myomiRs” [48] and have been associated with myogenesis and with various biological processes in skeletal muscle and/or heart [49–58]. In Nile tilapia, an increase in the expression of these three myomiRs is observed in muscle throughout the fish life [59]. The association between these miRNAs and active myogenesis thus appears to be evolutionarily conserved in vertebrates. A higher level of plasmatic myomiRs in well-fed animals compared to animals under a restricted diet would be consistent with the significant increase in growth rate observed in the present individuals when fed *ad libitum* [27]. Together, these observations suggest that plasma levels of miR-1-1/2-3p, miR-133-a-1/2-3p and miR-206-3p have the potential to identify episodes of active myogenesis. Under the hypothesis that these biomarker myomiRs would reflect muscle growth rate, which would require proper experimental validation using other samples and individuals in a variety of experimental situations, this result could offer a wide range of possible applications. For wild population management, these biomarkers could for instance offer the possibility to assess the quality of an ecosystem through the ability to monitor fish growth throughout the year. In aquaculture, it could allow fine phenotyping of muscle growth in response to specific diets or rearing conditions. More importantly, these biomarkers could allow to specifically question muscle growth in comparison to global body growth that can be influenced by the development of other tissues such as fat deposits, an information that is currently not easily accessible without sacrificing the fish. Finally, the identification of other biomarker c-miRNAs, such as markers of viral and bacterial infections, would allow the collection of a panel of relevant complementary information from a single blood sample and thus offer tremendous phenotyping possibilities.

**Figure 7:**
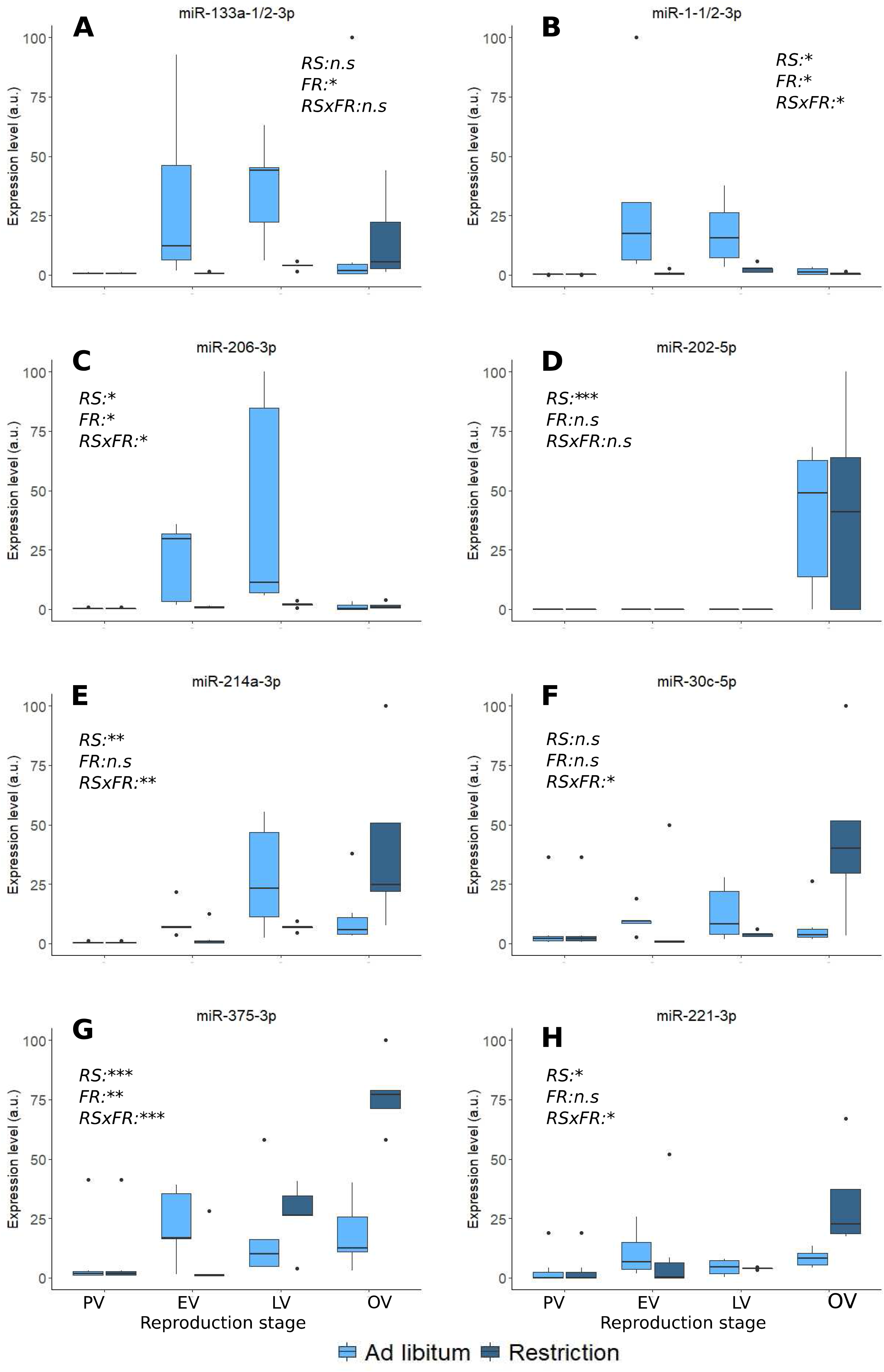
Plasma abundance of candidate biomarker miRNAs. Quantitative PCR analysis of selected miRNAs during the reproductive cycle and in response to feeding rate. The following reproductive stages were analyzed: pre-vitellogenesis (PV), early vitellogenesis (EV), late vitellogenesis (LV) and ovulation (OV). The feeding rate was either ad libitum (al) or restricted (r). Significant differences (ANOVA, p<0.05) are indicated by * for reproductive stage (RS), feeding rate (FR) and feeding rate – stage interactions (RS X FR). Replicates (N= 5) correspond to different individual fish.

**Figure 8:**
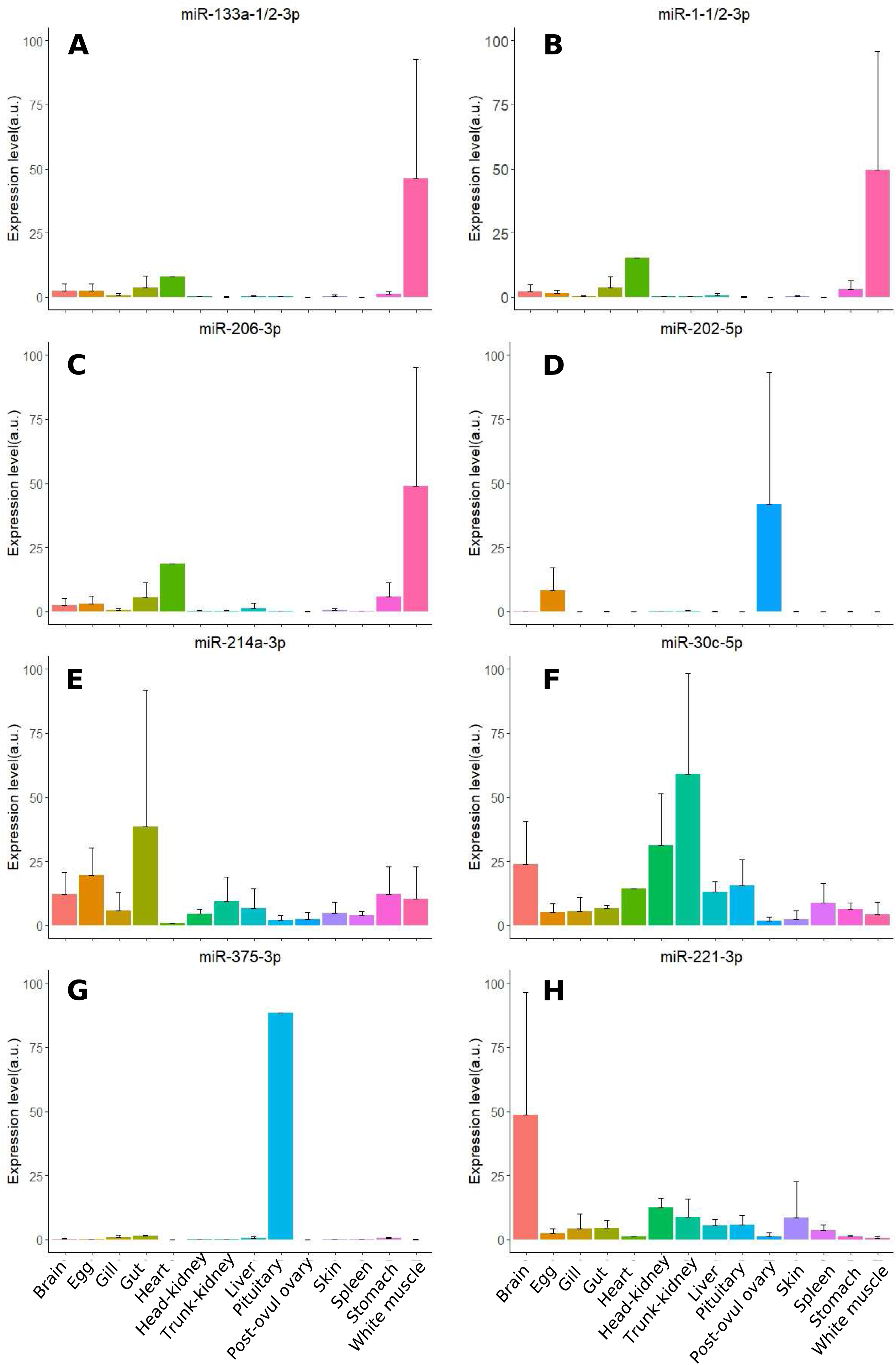
Organ abundance of candidate biomarker c-miRNAs. Quantitative PCR analysis of selected miRNAs in different organs. Replicates (N= 3) correspond to different individual fish. Means and SEM are displayed.

Among the miRNAs that we investigated to assess their potential use in non-invasive phenotyping, four c-miRNAs exhibited significant changes in their plasma abundance in response to feeding rate, either globally in the case of miR-375-3p (Figure 7G) or in interaction with the reproductive stage in the case of miR-214a-3p, miR-30c-3p, and miR-221-3p (Figure 7E-F, H). The latter, miR-214-a-3p, miR-30c-3p, and miR-221-3p, were also expressed in a wide variety of organs (Figure 8 E-F, H), making it hazardous to speculate on their organ of origin and the biological processes in which they may be involved. The expression profiles of these c-miRNAs, however, indicate that they could be used, most likely in combination with other specific c-miRNAs, to estimate the metabolic state of the fish at a given reproductive stage. Interestingly, these three c-miRNAs have been used as plasma biomarkers for several human pathologies such as cancer (liver, prostate, ovary and pancreas cancers) and cardiovascular and renal diseases [60–65]. In contrast, the highly predominant expression of miR-375-3p in the pituitary (Figure 8), which is consistent with existing data in other vertebrate species [66,67] and can tentatively be associated with the neuroendrocrine control of biological processes such as nutrition and reproduction. This c-miRNA exhibited highly significant differences in plasma abundance in response to feeding rate both globally and in a reproductive-stage-dependent manner (Figure 7). The difference in miR-375-3p levels in response to feeding rate was especially marked during late vitellogenesis and ovulation period. In other animal species, miR-375-3p has been implicated in the regulation of insulin [67] but also in the reproductive metabolism and a role in reproduction has been suggested [66,68]. Yuan et al. [66] identified the gonadotropin-releasing hormone (GnRH) receptor, a major player in the neuroendocrine control of reproductive cycles [69], as a predicted target gene of miR-375-3p in cattle. Yu and coworkers [68] showed that miR-375-3p is a key factor in regulating estradiol synthesis by mediating the CRH signaling pathway in pig. Even though the role of miR-375-3p in animal reproduction remains unclear, this miRNA appeared to be abundant in the plasma and highly responsive to changes in reproductive and metabolic states in female rainbow trout. For these reasons, miR-375-3p is a highly promising candidate biomarker for non-invasive phenotyping of neuroendocrine response in rainbow trout and possibly other animal species.

Among the c-miRNAs that we monitored by QPCR throughout the reproductive cycle, miR-202-5p had the most striking profile (Figure 7D). Independent of the feeding regime, miR-202-5p exhibited a dramatic increase in its plasma abundance at ovulation and was also among the most highly abundant c-miRNAs in ovarian fluid according to the small RNA-seq data. In fish, miR-202-5p plays a major role in reproduction and female medaka lacking expression of miR-202-5p produced fewer eggs and of lesser quality [40]. It is therefore possible that miR-202-5p in plasma and in ovarian fluid plays an important biological role at the time of ovulation that would require further investigations. As already described in rainbow trout, teleost fishes and other vertebrates, miR-202-5p is predominantly expressed in the ovary and was also detected in unfertilized eggs [28,40,70–72]. In the rainbow trout ovary, miR-202-5p is differentially regulated during oogenesis with a peak of expression during vitellogenesis followed by a stepwise decrease during final oocyte maturation [73]. The profile of miR202-5p in plasma reported here with a peak at ovulation thus differs from its ovarian expression. It is possible that this discrepancy results from the delay between expression in the ovary during vitellogenesis and accumulation in plasma during periovulatory period. This is, however, unlikely given the duration of several months between mid-vitellogenesis and ovulation. The sharp increase in plasma miR-202-5p levels at ovulation (Figure 7D), in contrast, suggests a release during the periovulatory period, either from the ovary or from the eggs. It is thus possible that a dynamic accumulation of miR-202-5p in plasma occurs either immediately prior to or following ovulation. Under this hypothesis, circulating miR-202-5p levels could serve as a biomarker to predict approaching ovulation, if the accumulation in the plasma occurs prior to ovulation, or to estimate post-ovulatory egg ageing, if the accumulation in the plasma occurs at or after ovulation. In both cases, this c-miRNA would be of major interest as a non-invasive phenotyping biomarker enabling, in aquaculture or wild resource management settings, the selection of females that are close to or at ovulation to prevent the occurrence of post-ovulatory ageing of the eggs, a phenomenon associated with a dramatic decrease in egg quality [74].

Together, both small RNA-seq and QPCR data revealed that the levels of selected circulating miRNAs exhibited major differences during the female rainbow trout reproductive cycle and, for some of them, also in response to changes in the metabolic state. Some of these c-miRNAs therefore appear to be highly relevant candidate biomarkers that could serve for non-invasive phenotyping of sexual maturation (i.e. progress into the reproductive cycle) and high episodes of muscle growth.

## 3 Conclusion

In the present study, we provide a reference rainbow trout miRNA repertoire annotation that was previously missing in this species and the first comprehensive analysis of plasma and ovarian fluid miRNAomes in a fish species. We show that biological fluid miRNAomes are extremely diverse and encompass a high proportion of the overall miRNAome of the species. While sharing common miRNAs, plasma and ovarian fluid miRNAomes nevertheless exhibited marked differences with fluid-specific combinations of highly abundant miRNAs and fluid-specific miRNAs. In addition, our data suggest that the complexity of c-miRNAomes in plasma and ovarian fluid originates, at least in part, from the accumulation of miRNAs expressed in a wide diversity of organs. Our results also raise the question of fluid-specific miRNAs that could result from a fluid-dependent accumulation of some c-miRNAs over time. We further showed that plasma exhibited major changes in c-miRNA abundances depending on the metabolic and the reproductive state. We subsequently identified a subset of three evolutionarily conserved myomiRs (miR-1-1/2-3p, miR-133-a-1/2-3p and miR-206-3p) that accumulated in plasma in response to high feeding levels and thus appear as strong candidate biomarkers of active myogenesis. We also identified miR-202-5p as a candidate biomarker of ovulation that could be used to predict ovulation and/or egg quality. Despite a lack of clear understanding of the biological roles of c-miRNAs, these highly promising results highlight the potential of c-miRNAs as physiologically relevant biomarkers and pave the way for the use of c-miRNAs for non-invasive phenotyping in many fish species.

## 4 Materials and Methods

### 4.1 Experimental design and fluid sampling

To generate contrasting physiological situation, two feeding strategies were used throughout the reproductive cycle in rainbow trout. Females were either fed *ad libitum* or 80% of ad libitum (restriction). These two feeding regimes were used to trigger contrasted metabolic states which proved to induce significant increases in fish weight and condition factors in fish fed *ad libitum* compared to fish under a restricted diet as previously described [27]. Plasma samples were obtained at the following stages of the reproductive cycle: previtellogenesis (PV), early-vitellogenesis (EV), late-vitellogenesis (LV) and ovulation (OV). Ovarian fluid was also sampled at ovulation. Reproductive stages were estimated based on existing data on rainbow trout reproductive cycle [75]. Ovulation was checked once a week and fish were sampled the next day. Ovulation was checked once a week and fish were sampled 2 days later. Ovulation (OV) stage thus corresponds to fish for which ovulated eggs have been present in the body cavity for 2-9 days. Blood samples were collected from the caudal vein using EDTA-treated syringes (sodium EDTA, 10%). Blood samples were centrifuged (3000g, 15 min, 4°C), and plasma samples were aliquoted, frozen in liquid nitrogen and stored at −80°C until further analysis. Ovarian fluid was collected after manual stripping of the eggs from ovulated females. Eggs were poured onto a mesh screen and ovarian fluid was collected, centrifuged (3000g, 15 min, 4°C) to remove cells and debris, frozen in liquid nitrogen and stored at −80°C until further analysis.

### 4.2 RNA preparation

For small-RNA sequencing (sRNA-seq), RNA extraction was carried out using pooled-samples of plasma and ovarian fluid. For both plasma and ovarian fluid, each pool was made from 50 individual that contributed equally. A total of 16 pooled-samples were used for sRNA-seq. For plasma, six experimental conditions were analyzed: three stages during oogenesis for each of the two feeding levels conditions (*ad libitum* vs restriction). For each condition, two pools were analyzed resulting in a total of 12 small RNA libraries. For ovarian fluid, duplicates for each feeding levels were sequenced resulting in a total of four small RNA libraries.

For quantitative PCR (QPCR) validation, extractions were carried out on individual plasma samples. RNA was extracted from five samples per condition and a total of ten conditions were analyzed. Validation by QPCR was carried out only on plasma samples. Samples were randomly taken from all four time-points during oogenesis and for each of the two feeding conditions (*ad libitum* vs restriction).

For both small RNA-seq and QPCR, samples were homogenized in Trizol reagent (Macherey-Nagel) at a ratio of 400 μL of fluid per milliliter of reagent and total RNA was extracted according to manufacturer’s instructions. During the RNA extraction protocol, glycogen was added to each sample to facilitate visualization of precipitated RNAs. Expression of selected candidate miRNAs was analyzed by QPCR in 14 different organs and tissues (brain, pituitary, gills, head kidney, trunk kidney, spleen, liver, stomach, intestine, post-ovulatory ovary, egg, skin, heart, and white muscle) that were sampled from three ovulated females. Tissues were homogenized in Trizol reagent at a ratio of 100mg per milliliter of reagents and total RNA was extracted according to the manufacturer’s instructions.

### 4.3 Small RNA sequencing

Libraries were constructed using the NEXTflex small RNA kit v3 (Bioo Scientific). Starting from 1μg of total RNA, an adapter was ligated on the 3’ end of the RNAs. A second adapter was ligated to the 5’ end. Ligated RNAs were subjected to reverse transcription using M-MuLV transcriptase and a RT primer complementary to the 3’ adapter. PCR amplification (16 cycles) was performed on the cDNA using a universal primer and a barcoded primer. Final size selection was performed on 3% gel cassette on a Pippin HT between 126pb and 169pb. Sequencing (single read 50 nucleotides) was performed using a HiSeq2500 (Illumina) with SBS (Sequence By Synthesis) technique. A total of over 187 million reads (after Illumina Purity Filter) were obtained with a number of read per library ranging from 12.1 to 15.0 millions. Raw reads were deposited into NCBI Sequence Read Archive under accession number PRJNA631932. Reads were trimmed of the adaptor sequence GCCTTGGCACCCGAGAATTCCA and of the random primers using Cutadapt [76].

### 4.4 Establishment of a reference rainbow trout miRNAome

The rainbow trout miRNAome annotation was established using Prost ![31], which was first run using the rainbow trout reference genome (NCBI RefSeq assembly accession GCF_002163495.1) and all ovarian fluid and plasma reads generated in the present study. This was performed using a set of five manually curated fish mature and hairpin sequences (*Lepisosteus oculatus, Danio rero*, Poecilia Mexicana, *Gasterosteus aculeatus*, and *Chaenocephalus aceratus*) [30,32,33]. *Prost!* was run using the zebrafish as focal species and with proposed default settings (minimum read count of 20, sequence size comprised between 17 and 25). The resulting set of annotated genomic locations was then curated using the recommendations of *Prost!* documentation. First, reads with at least one mismatched nucleotide to the genome, or aligning to more than 20 locations, were not considered for annotation. Then, homology to previously described miRNAomes was used to build a set of annotated mature miRNA sequences in rainbow trout. When miRNAs displayed sequence variations aligning to the same locus or loci (i.e., isomiRs) [77], the most abundant isomiR for the locus was the one annotated. Then, homology to previously described miRNAomes [29–31] was used to build a set of annotated rainbow trout mature miRNA sequences. When miRNAs displayed sequence variations aligning to the same locus or loci (i.e., isomiRs), the most abundant isomiR for the locus was the one annotated.

To provide a comprehensive rainbow trout miRNAome annotation, this set was further extended by rerunning Prost! with the same settings but on samples originating from a wide variety of tissues, organs and cell types [28] and using a combination of the above described rainbow trout annotation and all existing teleost miRNA mature and hairpin sequences, with comparable settings and the same curation process. The resulting rainbow trout miRNAome annotation is freely available here: https://github.com/INRAE-LPGP/microRNA.

### 4.5 *miRNA* quantification and differential expression analysis

Prost! was run using the newly obtained rainbow trout miRNA annotation, all the circulating fluids sequencing samples, and identical settings as for the annotation step. Normalized miRNA counts in Reads Per Million (RPM) were extracted from the “compressed by annotation” Prost! output tab. When Prost! identified multiple potential annotations for a given sequence, counts for this sequence were distributed evenly between the annotations. Counts in samples from the same tissue were averaged, and miRNAs for which normalized abundance was greater than 10 RPM were considered expressed in a given tissue. Raw and normalized read counts are provided in additional file 4.

To evaluate putative organ origin of circulating miRNAs, we associated each c-miRNA to the organ in which it was the more abundant, under the hypothesis that it represents the organ from which it was the most likely to originate, or at least one the major contributors. For this analysis we excluded several types of samples, including eggs, spermatogonia, testis, whole embryo, myoblasts, myotubes, and leucocytes. Each c-miRNA was therefore assigned to one of 13 organs (brain, gills, heart, head kidney, intestine, liver, muscle, ovary, pituitary, skin, spleen, stomach, trunk kidney).

Expression data across organs were visualized using a heatmap based on the expression matrix centered and reduced by row and generated with the R package *heatmaply* [78]. Expression within the 14 fluid samples was visualized by performing a Principal Component Analyses (PCA) on log-transformed DESeq2 counts. To account for the greater importance of abundant miRNAs, the PCAs were centered but not scaled. PCAs were computed using the R package *FactoMineR* [79] and 95% confidence ellipses associated to the different condition were drawn using the plotellipses function. In the case of previtellogenesis plasma samples, only two samples were available, and no ellipse could therefore be drawn.

Differential expression analyses were performed using DESeq2 [80] and raw counts from all 14 fluid miRNA expression data. For each differential expression test, log fold changes were corrected using lfcShrink(type=”apeglm”), and p-values were corrected using the FDR method to account for multiple testing.

To identify miRNAs differentially expressed between plasma and ovarian fluid samples, a model was built using all samples, and considering all possible effects (~Sample + Feeding + Time). Differential expression was tested using a Plasma vs. Ovarian fluid contrast (i.e., “Sample” contrast). Differential expression of miRNAs between *ad libitum* and restricted feeding was further tested independently in plasma and ovarian fluid samples with the same parameters.

Differential expression across the three considered reproductive stages was evaluated only for the 10 plasma samples because ovarian fluid was sampled only at ovulation. Differential expression was tested using a likelihood ratio test between the “~ Time + Feeding” model and the “~Feeding” model. Afterwards, expression trajectories for the differentially expressed miRNAs were clustered using the function degPatterns of the R package *DEGReport* [81] on a regularized log transformed DESeq2 count matrix.

### 4.6 QPCR validation of miRNA expression and statistical analysis

Expression of circulating and organ microRNAs was assessed using the TaqMan Advanced miRNA Assay (A25576, Applied Biosystems, ThermoFisher) according to manufacturer’s instructions. Synthetic cel-miR-39 mimic (miScript miRNA Mimics, QIAGEN) was spiked to each RNA sample at a ratio of 11 fmol per ug of total RNA and used as an exogenous control for normalization. Briefly, 1.75 ng of total mi-RNA were used for the initial poly(A) tailing, ligation and reverse transcription reactions to synthesize the cDNA of all miRNAs followed by a pre-amplification step. RT-QPCR assays were performed on the Roche LightCycler 480 system (Roche Diagnostics, France). The assays were carried out in a reaction mix of 10 μL containing 2.5 μL of cDNA (diluted 5 times), 5 μL of 2X Fast Advanced Master mix (Applied Biosystems, USA), 0.5 μL of TaqMan Advanced miRNA Assay (20X) (Applied Biosystems, USA) and 2 μL of DNAse/RNAse free water. Quantitative RT-PCR was performed using the LightCycler 480 System (Roche Life Science) with the following conditions: 95°C for 20 sec; and 40 cycles of 95°C for 1 sec and 60°C for 20 sec. The relative expression of miRNA within a sample set was calculated from a standard curve using LightCycler 480 System software release 1.5.1.62. All RT-QPCR were performed in duplicates. Data was normalized using cel-miR-39 expression levels. Sequences of the miRNA probes are provided in additional file 5. Expression levels measured by RT-QPCR in Figures 7 and 8 were given as means and standard deviations across biological replicates. Statistical analysis of the data was carried out using R studio software. Two-way ANOVA was performed to assess the effect of feeding level and reproductive stage on miRNA abundance.

## 5 Declarations

### 5.1 Ethics approval

Experiments were conducted at INRAE PEIMA experimental facilities (Sizun, France) as previously described [27]. Experiments and procedures were fully compliant with French and European animal welfare policies and followed guidelines of the INRAE PEIMA Institutional Animal Care and Use Ethical Committee, which specifically approved this study.

### 5.2 Availability of data and materials

The datasets generated and analyzed during the present study are available in NCBI SRA repository, https://www.ncbi.nlm.nih.gov/sra/PRJNA631932 (rainbow trout biological fluids), https://www.ncbi.nlm.nih.gov/bioproject/PRJNA227065 (rainbow trout tissues, organs, cell types and embryos).

### 5.4 Authors’ contributions

EC performed RNA extractions, QPCR analyses, participated in miRNA annotation and in data analysis and drafted the manuscript. CG performed small RNA-seq analyses including statistical analyses, participated in miRNA annotation and in manuscript preparation, TD participated in miRNA annotation; in data analysis, and in manuscript preparation, JM participated in small RNA-seq processing and analysis, SG performed small RNA sequencing, JHP participated in data analysis and in manuscript preparation, SS co-conceived the study, JB co-conceived the study, participated in miRNA annotation and in data analysis, and prepared the manuscript. All authors read and approved the final manuscript.

### 5.5 Funding

This work was supported by France Génomique National infrastructure, funded as part of “Investissement d’avenir” program managed by Agence Nationale pour la Recherche (contract ANR-10-INBS-09), by European Commission/European Fund of Maritime Affairs and Fisheries (NutriEgg, PhenomiR), and by NSF USA grant PLR-1543383, NIH USA grant R01 OD011116.

#### 5.6 Acknowledgements

The authors thank INRAE PEIMA staff for fish rearing, MGX staff for small RNA-sequencing and GenOuest facility for providing computing infrastructures.

## 7 Additional Files

### Additional file 1

Average normalized read counts (RPM) for all annotated miRNAs detected in biological fluids and organs.

### Additional file 2

Differentially expressed miRNAs between ovarian fluid and plasma samples.

### Additional file 3

Differentially expressed miRNAs in plasma during reproductive cycle or in response to feeding rate.

### Additional file 4

Normalized and raw read counts in all analyzed libraries

### Additional file 5

QPCR primer sequences

